# Engram Reactivation Mimics Cellular Signatures of Fear

**DOI:** 10.1101/2023.12.14.571734

**Authors:** Rebecca L. Suthard, Ryan A. Senne, Michelle D. Buzharsky, Anh H. Diep, Angela Y. Pyo, Steve Ramirez

**Affiliations:** Graduate Program for Neuroscience, Boston University, Boston, MA, 02215, USA; Department of Biomedical Engineering, Boston University, Boston, MA, 02215, USA; Department of Psychological and Brain Sciences, The Center for Systems Neuroscience, Neurophotonics Center, and Photonics Center, Boston University, Boston, MA, 02215, USA

**Keywords:** Hippocampus, engram, astrocyte, fiber photometry, optogenetics, fear, memory

## Abstract

Engrams, or the physical substrate of memory in the brain, recruit heterogeneous cell-types. Targeted reactivation of neurons processing discrete memories drives the behavioral expression of memory, though the underlying landscape of recruited cells and their real-time responses remain elusive. To understand how artificial stimulation of fear affects intra-hippocampal neuronal and astrocytic dynamics as well as their behavioral consequences, we expressed channelrhodopsin-2 in an activity-dependent manner in dentate gyrus neurons while performing fiber photometry of both cell types in ventral CA1 across learning and memory. Neurons and astrocytes were shock-responsive, while astrocytic calcium events were uniquely modulated by fear conditioning. Notably, optogenetic stimulation of a hippocampus-mediated engram recapitulated coordinated calcium signatures time-locked to freezing that were also observed during natural fear memory recall, suggesting that engram activation alters activity across different cell types within hippocampal circuits during the behavioral expression of fear. Together, our data reveals cell-type specific hippocampal dynamics during freezing behavior and points to neuronal-astrocytic coupling as a shared mechanism enabling the natural and artificial recall of a memory.

**Highlights:**

- Ventral hippocampal neurons and astrocytes are active during foot shock
- Calcium activity is time-locked to freezing during fear conditioning and recall
- Optogenetic reactivation of fear recapitulates cellular signatures seen during recall
- Reactivation of a fear memory allows prediction of freezing behavior

## Introduction

The hippocampus (HPC) is a region within the medial temporal lobe that is indispensable for episodic memory ^1^. In recent years, there has been an increasing focus on the ventral HPC’s vital role in emotional processing and memory. Studies have confirmed differential roles along the dorsal and ventral axis, where the dorsal hippocampus (dHPC) is necessary for spatial and temporal encoding, while lesions to the ventral hippocampus (vHPC) cause emotional dysregulation and impaired stress responses ^2–4^. More specifically, the ventral CA1 (vCA1) subregion contains a subset of basolateral amygdala-projecting neurons necessary for contextual fear conditioning (CFC) and are responsive to the aversive conditioning^3^. This body of work suggests a critical role for vCA1 in aversive conditioning and the subsequent emotional responses.

The hippocampus’s subregions and cell types have a complex and heterogeneous structure-function relationship. The dentate gyrus (DG), CA3, and CA1 subregions of the hippocampus make up the “trisynaptic loop,” which has shown to be a critical circuit for HPC mnemonic processes. Moreover, the dHPC, specifically the DG, contains defined sets of cells that undergo plasticity-related changes during learning which, when activated, are sufficient to drive the behavioral expression of memory ^5–9^. These studies rely on inducible genetic tools, such as the Tet-tag system^10^, to effectively ‘tag’ or label neuronal ensembles or *engrams* active during an initial experience which can later be reactivated via chemo- or optogenetics. This field of work has enhanced our understanding of HPC computations and allows for a time-locked strategy for assessing a causal role between circuitry and behavior. However, most studies to date have focused almost exclusively on neuronal contributions to memory engrams, while the role of non-neuronal cell types within this hippocampal circuitry have remained relatively understudied. Thus, it is unknown whether these cells modulate intrahippocampal population dynamics in a pattern akin to natural memory recall.

Astrocytes are a predominant glial cell type that have been shown to play a pivotal role in the tripartite synapse, coupling with pre- and postsynaptic neurons to bidirectionally modulate synaptic communication ^11–15^. These cells perform vital functions including maintaining the blood-brain barrier, supporting metabolic needs of surrounding neurons, releasing gliotransmitters and expressing neurotransmitter receptors to appropriately respond to and modulate their neuronal neighbors ^16–18^. Recently, there has been a shift to studying astrocytes at the systems-level through the use of optogenetics, chemogenetics and optical imaging techniques ^19–23^. For instance, manipulation of astrocytes in hippocampal and amygdalar sub-regions has been shown to impair or enhance both recent and remote memory formation and retrieval ^24–27^. Moreover, recent engram studies have demonstrated the importance of non-neuronal cell types, microglia, and oligodendrocytes, in experience- and activity-dependent processes ^28–30^. Broadly, this burgeoning field suggests that glial cells are playing a role in modulating local and long-range projections within the brain to regulate behavior. Still, despite these recent advances little is known about how astrocytes may be playing a role in emotional processing within vCA1 during fear memory expression.

To that end, we combined activity-dependent tagging strategies with dual-color fiber photometry to study neuronal-astrocytic interplay of activity across fear encoding, natural recall, and artificial reactivation of a tagged engram. We injected a glial fibrillary acidic protein (GFAP)-driven GCaMP6f virus paired with a neuronal human synapsin (hSyn)-driven jRGECO1a in ventral hippocampal CA1 (vCA1) to measure neuronal and astrocytic dynamics, while concurrently tagging and reactivating cells active during contextual fear conditioning (CFC) in the dorsal dentate gyrus (dDG) using Tet-tag viral constructs. We find that optogenetic reactivation of a fear memory is sufficient to induce cellular signatures that are akin to those seen during natural recall. These findings further enhance our understanding of the complex interplay of the cellular machinery of the brain and suggest that neurons and astrocytes are important members of the same processes.

## Results

### In vivo calcium recordings of neurons and astrocytes in the ventral hippocampus during natural and artificial memory reactivation

We first monitored neuron and astrocyte calcium dynamics in hippocampal vHPC using dual-color fiber photometry in freely moving mice across habituation, CFC, natural recall and optogenetic (‘artificial’) reactivation of a tagged fear memory. Wild type mice were injected in vCA1 unilaterally with AAV5- GfaABC1D-GCaMP6f and AAV9-hSyn-jRGECO1a to express genetically-encoded calcium indicators (GECIs) in astrocytes and neurons, respectively. Additionally, these mice were bilaterally injected with the Tet-tag viral cocktail of AAV9-c-fos-tTA and AAV9-TRE-ChR2-eYFP or eYFP control in the dorsal DG to allow for temporal control over ‘tagging’ and reactivating a neuronal ensemble with this simultaneous recording of calcium dynamics (Figure 1A). jRGECO1a and GCaMP6f successfully and effectively expressed in vHPC neurons and astrocytes, respectively (Figure 1B). Across all days of behavior, we used fiber photometry to record freely moving neuronal and astrocytic calcium activity in the vHPC (Figure 1C). Mice were placed into three groups: Shock-ChR2 (mice that received foot shock and expressed active channelrhodopsin (ChR2) in tagged fear engram cells), Shock-eYFP (mice that received foot shock and expressed control eYFP in tagged fear engram cells) and Neutral-ChR2 (mice that did not receive foot shock and express active ChR2 in tagged neutral engram cells). On Day 1, all mice remained on doxycycline (Dox) diet while they freely explored neutral Context B (Cxt B) and received blue-light stimulation for habituation. After this session, mice had Dox diet replaced with standard chow for 48 hours prior to the engram labeling. On Day 3, mice in the Shock-ChR2 and Shock-eYFP groups underwent CFC in Context A (Cxt A). Here, they received 4, 0.75mA foot shocks while off Dox. Mice in the Neutral-ChR2 group were placed in Cxt A for the same duration of time in the absence of aversive stimuli. After this session, all mice were returned to the Dox diet to close the ‘tagging window’. On Day 4, all mice were placed back in Cxt A for contextual (‘natural’) recall. On Day 5, all mice were placed in the previously habituated Cxt B where they received blue-light stimulation during the 120-240 seconds and 360-480 second time periods. 90 minutes after the first light-ON epoch of optogenetic stimulation, mice were transcardially perfused to capture peak cFos protein levels and allow us to examine the ‘reactivation rate’ of tagged vs. reactivated neurons in dDG (Figure 1C).

**Figure 1.**
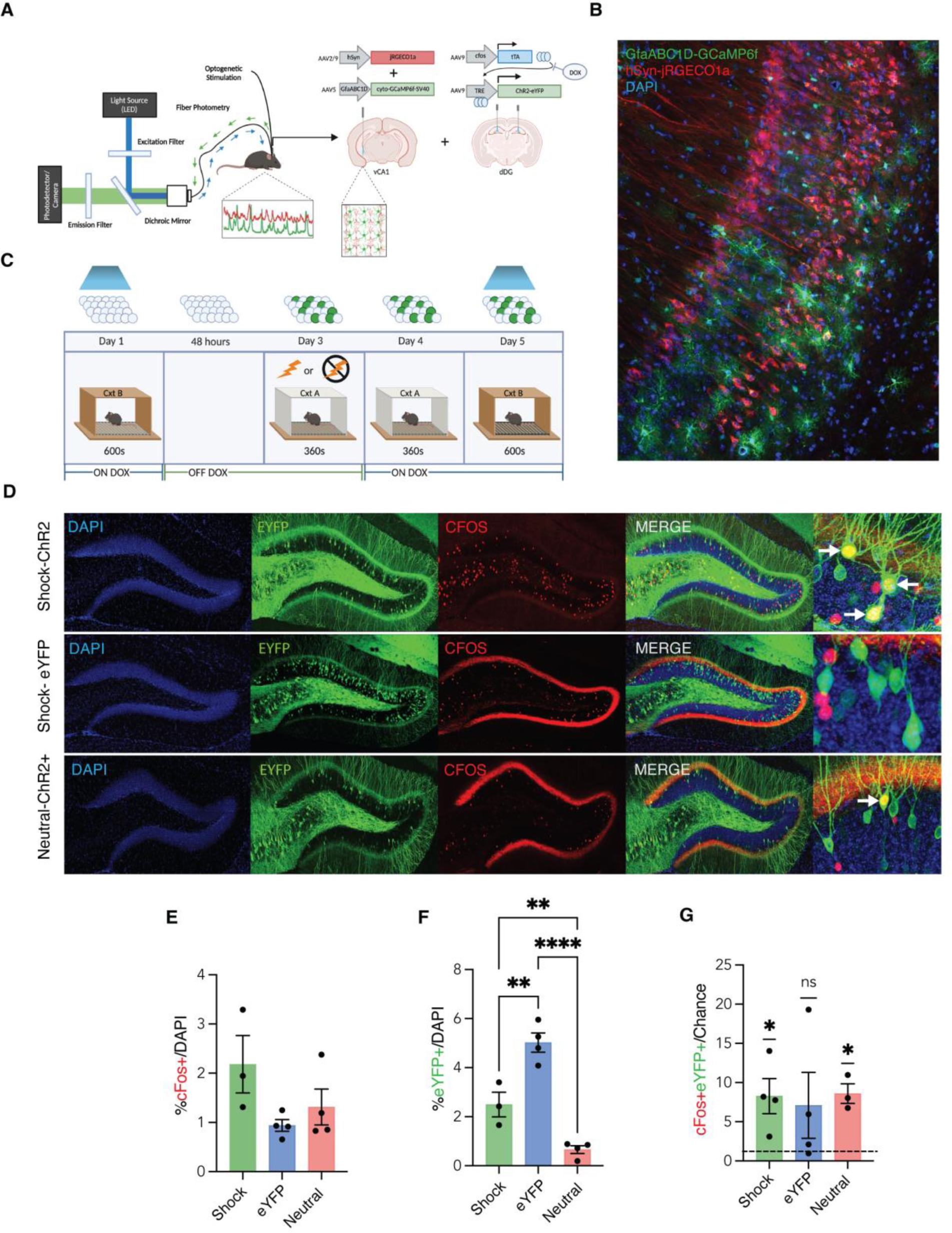
In vivo calcium recordings of neurons and astrocytes in the ventral hippocampus during natural and artificial memory reactivation. (A) Neuron-astrocyte fiber photometry recordings hippocampal ventral CA1 (vCA1) coupled with optogenetic stimulation of a dorsal dentate gyrus (dDG)- mediated neuronal ensemble. Mice were injected in vCA1 with AAV5-GfaABC1D-cyto-GCaMP6f-SV40 and AAV9-hSyn-jRGECO1a-WPRE.SV40, and received a unilateral optical fiber implant to enable Ca2+ recordings. dDG was infused with AAV9-c-fos-tTa and AAV9-TRE-ChR2-eYFP or control vector, and implanted with bilateral optical fibers. (B) Representative confocal microscopy image of the vCA1 pyramidal cell layer expression of hSyn-jRGECO1a (red; neurons), GfaABC1D-GCaMP6f (green; astrocytes) and DAPI (blue; nuclei). (C) Schematic representation of the viral tagging strategy used during behavioral testing: On Day 1, mice were placed in Context B (Cxt B) for 600 seconds of habituation in the presence of blue light optogenetic stimulation while on DOX diet. DOX diet was removed for 48 hours immediately after this session to open the ‘tagging’ window. On Day 3, mice were split into three groups: Shock-ChR2, Shock-eYFP and Neutral-ChR2, where they underwent a neutral (no-shock) or contextual fear conditioning (4-shock; 0.75mA, 2 second duration foot shocks) experience for 360 seconds in Context A (Cxt A). Mice were placed immediately back on DOX diet to close the ‘tagging’ window. On Day 4, all mice were placed back in Cxt A for a 360 second natural recall session. Day 5 mice underwent a 600 second session in Cxt B where they received optogenetic blue light stimulation (450nm laser diode, 20Hz, 10ms pulse, 15mW) to ‘reactivate’ the tagged ensemble. 90 minutes after the start of optogenetic stimulation, mice were transcardially perfused to capture peak endogenous cFos protein resulting from stimulation. (D) Representative 20x confocal images of dDG histology visualizing DAPI+ cells (blue), eYFP+ cells (green; active during FC), cFos+ cells (red; active during optogenetic stimulation) and overlaps (yellow; active during both sessions) between these three channels for each group: shock-ChR2 (top), shock-eYFP (middle) and neutral-ChR2 (bottom). Zoomed representative 20x images of these overlaps (yellow) are shown in the far-right of each row with arrows highlighting these. (E) %cFos+/DAPI+ cells were not different across groups; One-way ANOVA. (F) %eYFP+/DAPI+ cells was significantly increased in Shock-ChR2 and Shock-eYFP groups; One-way ANOVA and Holm-Sidak’s multiple comparisons. (G) %cFos+eYFP+/chance (overlap) cells were significantly greater than chance in Shock-ChR2 and Neutral-ChR2 groups; One-sample t-tests of each group against chance (1.0). Error bars indicate SEM. p ≤ 0.05, **p ≤ 0.01, ***p ≤ 0.001, ****p ≤ 0.0001, ns = not significant. Per group: n=3-4 mice x 8 dDG regions of interest (ROI) each were quantified for statistical analysis of cell counts.

Next, we performed histology to visualize the tagged dDG engram cells (green), reactivated cFos cells (red), DAPI (blue) and overlaps (yellow) for Shock-ChR2 (Figure 1D; top), Shock-eYFP (Figure 1D; middle) and Neutral-ChR2 (Figure 1D; bottom). There were no statistically significant differences in the active cFos+ cells (%cFos+/DAPI+) across any group (Figure 1E). The percentage of tagged cells (%eYFP+/DAPI+) was significantly different across groups. *Post-hoc* multiple comparisons demonstrated significantly greater eYFP+ ‘tagged’ engram cells in the Shock-eYFP group compared to both the Shock-ChR2 and Neutral-ChR2 groups (Figure 1F). Additionally, there was a significant increase in eYFP+ tagged cells in the Shock-ChR2 compared to the Neutral-ChR2 group (Figure 1F). Finally, to assess the similarity of the initially ‘tagged’ engram cells and those reactivated during optogenetic stimulation, we quantified the number of overlaps for each group cFos+eYFP+/DAPI+ compared to chance (eYFP+/DAPI+) x (cFos+/DAPI+). We observed that the overlaps/chance in the Shock-ChR2 and Neutral-ChR2 groups were significantly greater than the chance theoretical mean of 1.0 (Figure 1G). This result indicates that optogenetic stimulation of ‘active’ ChR2 protein in our tagged cells, both fearful and neutral in nature, increased the number of reactivated cells from the initially labeled experience, whereas our eYFP group was not activated.

### Neurons and astrocytes respond robustly to foot shock and freezing epochs during contextual fear conditioning

To test the hypothesis that astrocytes and neurons play an active role in the acquisition of contextual fear, we used *in vivo* fiber photometry to record the activity of both cell types across all experimental days of our behavioral task (Figure 1A; bottom). During CFC, mice in the Shock-ChR2 and Shock-eYFP groups had significantly increased freezing levels compared to Neutral-ChR2 mice that did not receive foot shock (Figure 2A-B). These mice acquired fear across the session, with an increase in the Shock-ChR2 and Shock-eYFP groups, but not in the Neutral-ChR2 (Figure 2A-B). Calcium timeseries for astrocytes (Figure 2C) and neurons (Figure 2D) in the Shock-ChR2 and Shock-eYFP groups indicated increased activity at the times of 0.75mA foot shock (120, 180, 240, 300 seconds), but not in the Neutral-ChR2 group that did not receive foot shock. To further understand this assessment, whole session peri-event analysis for astrocytes (Figure 2E) and neurons (Figure 2J) demonstrated a significant increase in activity at the onset of each foot shock, indicated by the dashed vertical lines. During the foot shocks, we observed a significant increase in the z-scored %dF/F for both astrocytes (Figure 2F) and neurons (Figure 2K) in the Shock-ChR2 and -eYFP groups, but not in the Neutral-ChR2 group, as expected. Further, calcium event characteristics were calculated for neurons and astrocytes in all groups. Astrocytes that underwent shock displayed an increase in peak height (%dF/F) and area under the curve (AUC), as well as a decrease in full-width half-maximum (FWHM) compared to those that did not receive shock (Figure 2G-I). On the other hand, neuronal calcium characteristics were not significantly different across any groups (Figure 2L-N).

**Figure 2:**
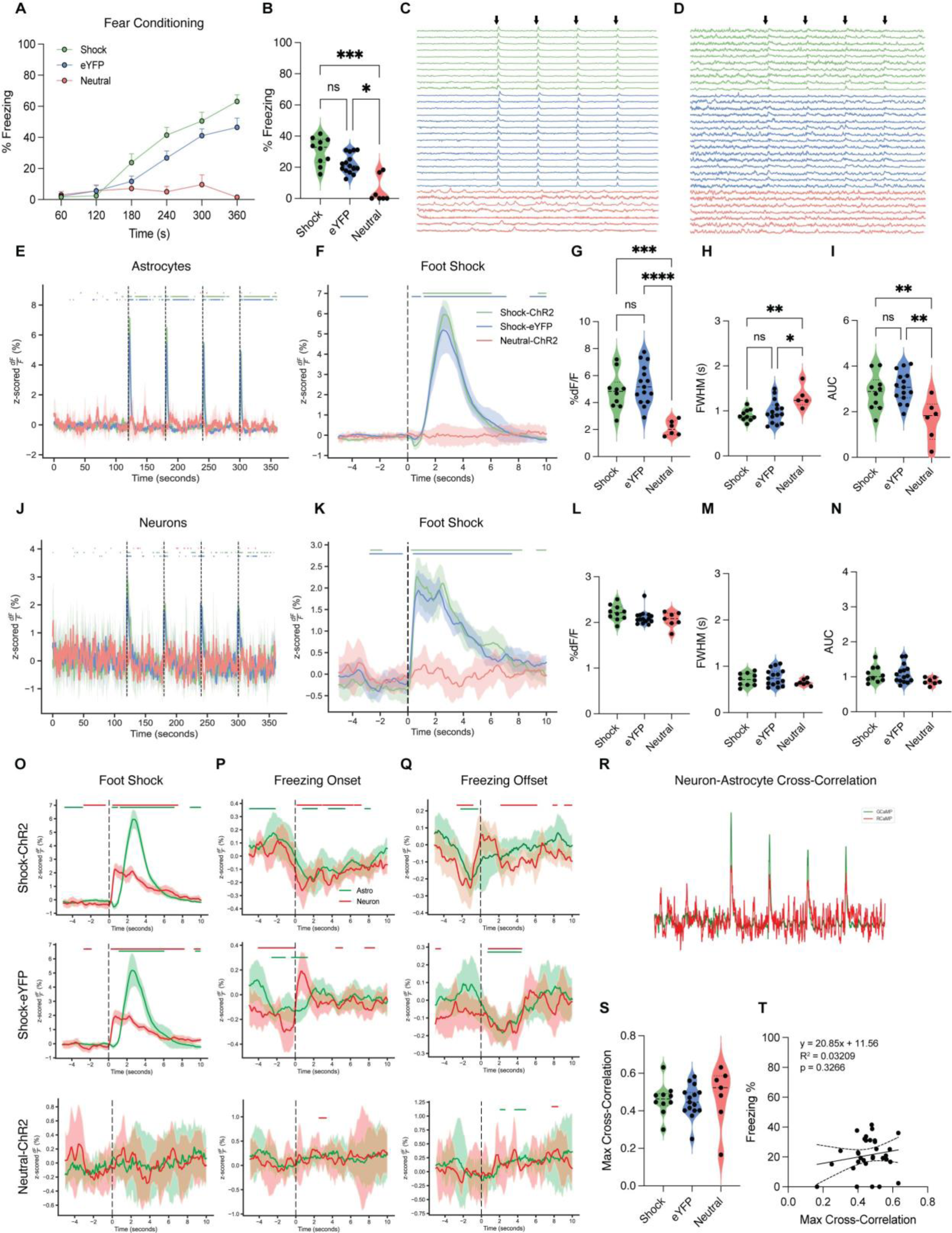
Neurons and astrocytes respond robustly to foot shock and freezing epochs during contextual fear conditioning. (A-B) Mice in Shock-ChR2 (green) and Shock-eYFP (blue) groups acquired fear during contextual fear conditioning (CFC) as shown by increased freezing percentage across the session (A) and on average (B) compared to Neutral-ChR2 (coral) mice that did not receive foot shock. (C-D) Calcium timeseries for astrocytes (C) and neurons (D) in the Shock-ChR2 and Shock-eYFP groups indicated increased activity at the times of 0.75mA foot shock (black arrows; 120, 180, 240, 300 seconds), but not the Neutral-ChR2 group. Each row represents a single subject across time (seconds). (E, J) Whole session peri-event analysis for astrocytic (E) and neuronal (J) calcium activity during CFC show an increase in activity (z-scored %dF/F) at the onset of each foot shock (dashed lines) in the Shock-ChR2 and Shock-eYFP groups. (F, K) Foot shock peri-event analysis for astrocytes (F) and neurons (K), with the onset of shock occurring at the dashed line (time = 0). Shock-ChR2 (green) and shock-eYFP (blue) show a significant increase in calcium activity (z-scored %dF/F), but neutral-ChR2 (coral) mice do not. (G-I, L-N) Calcium event metrics (z-scored) for astrocytes (G-I) and neurons (L-N); (G,L) peak height (%dF/F), (H,M) full-width half maximum (s), and (I,N) area under the curve (AUC). Astrocytes in Shock-ChR2 and Shock-eYFP groups show increased peak height and AUC, and decreased FWHM compared to Neutral-ChR2. Neurons show no significant differences in any metric during CFC across groups. (O-Q) Peri-event analysis for neurons (red) and astrocytes (green) during foot shock (O), freezing onset (P) and freezing offset (Q) for Shock-ChR2 (top), Shock-eYFP (middle) and Neutral-ChR2 (bottom). Shock-ChR2 and Shock-eYFP mice show a significant response across cell types to foot shock, freezing onset and offset, compared to Neutral-ChR2 mice. (R) Representative neuron-astrocyte cross-correlation, with neurons (red) and astrocytes (green) shifted to maximally align with one another. (S) Maximum cross-correlation value (0.0-1.0) was not significantly different across groups during CFC. (T) Simple linear regression for average freezing percentage (%) and maximum cross-correlation value for all mice collapsed across groups during CFC did not show a significant relationship. For all violin plots, One-way ANOVA or Kruskal-Wallis tests were performed on normal and nonparametric data, respectively. For post-hoc multiple comparisons tests, Holm-Sidak’s (normal) or Dunn’s (nonparametric) multiple comparisons were performed with p ≤ 0.05, **p ≤ 0.01, ***p ≤ 0.001, ****p ≤ 0.0001, ns = not significant. For event metrics and freezing behavior, shock n=10, eYFP n=15, neutral n=6-7; only one mouse in the neutral group was excluded as an outlier for astrocytic FWHM. For foot shock peri-event analysis, shock n=10, eYFP n=15, neutral n=7. For freezing onset and offset peri-events, shock n=10, eYFP n=15, neutral n=3; four neutral mice were excluded due to their entire lack of freezing during CFC.

Next, we hypothesized that astrocytes and neurons would be time-locked to the onset and offset of behavioral freezing bouts, in line with previous work from basolateral amygdala astrocytes during fear learning ^31^. Peri-event analysis of foot shock showed that neurons and astrocytes are both time-locked to 0.75mA foot shock, with neurons displaying a smaller amplitude event that begins shortly before astrocytic calcium (Figure 2O; top and middle), compared to those that did not receive an aversive stimulus (Figure 2O; bottom). For freezing onset, both neurons and astrocytes displayed a significant decrease in activity immediately before the start of the freezing bout, rebounding around 1-2 seconds post-event (Figure 2P; top and middle) that we did not observe in Neutral-ChR2 mice (Figure 2P; bottom). Interestingly, both astrocytes and neurons displayed a significant decrease in activity at the offset of freezing bouts in the Shock-ChR2 and -eYFP groups with little delay between the two cell types (Figure 2Q; top and middle). As neurons and astrocytes showed coordinated activity at freezing bouts, we hypothesized that as freezing percentage increased, there would be an increase in correlation between cell types (max cross-correlation). To understand this, we performed cross-correlation between the neuron and astrocyte calcium timeseries for the entire CFC session (Figure 2R). We observed that the maximum cross-correlation value was not significantly different across the three groups (Figure 2S). Further, we performed a simple linear regression to see if an increase in maximum neuron-astrocyte cross-correlation can predict an increase in an animal’s freezing level. For CFC, we did not observe a significant relationship between cross-correlation and average percent freezing (Figure 2T).

### Neuronal and astrocytic calcium are responsive to freezing epochs during contextual recall

To understand vHPC neuron-astrocyte calcium dynamics during natural recall, mice were placed back in CxtA in the absence of any aversive stimuli the next day. During recall, mice in the Shock-ChR2 and Shock-eYFP groups expressed high levels of fear, as shown by increased freezing across the session (Figure 3A-B) compared to Neutral-ChR2 mice that did not experience foot shock the day before. Whole session individual calcium timeseries (Figure 3C-D) and average traces (Figure 3E-F) for astrocytes and neurons showed qualitatively similar activity in shock groups during recall. Similar to CFC, there was a significant dip and rapid increase in the z-scored %dF/F for both astrocytes and neurons at the onset of freezing bouts for Shock-ChR2 and Shock-eYFP groups (Figure 3G; top and middle) that was not observed in the Neutral-ChR2 calcium (Figure 3G; bottom). Further, we observed a similar rapid decrease in calcium activity across cell types at the offsets of freezing bouts (Figure 3H; top and middle), that again was not observed in the Neutral-ChR2 mice (Figure 3H; bottom). Calcium event characteristics were calculated for both neurons and astrocytes across all groups during recall. Similar to CFC, astrocytes in the Shock-ChR2 group displayed significantly increased peak height and AUC compared to the Neutral-ChR2 group (Figure 3I). The Shock-eYFP group had a trend towards the same significant increase in peak height and AUC, with no difference between Shock-ChR2 and Shock-eYFP (Figure 3I). Again, neuronal calcium events did not differ significantly in any metric across groups during this session (Figure 3L-N). We further investigated the relationship between neuron-astrocyte cross-correlation and freezing during recall, as consolidation of the memory may contribute to a stronger relationship between these factors. We observed that the maximum cross-correlation value was significantly higher in the groups that received foot shock previously compared to Neutral-ChR2 (Figure 3O). Finally, a simple linear regression revealed a significant positive relationship between the maximum cross-correlation and freezing with an increase in neuron-astrocyte correlation predictive of an increase in the animal’s freezing behavior. With this information, we can conclude that increased maximum cross-correlation is positively associated with freezing levels (Figure 3P).

**Figure 3:**
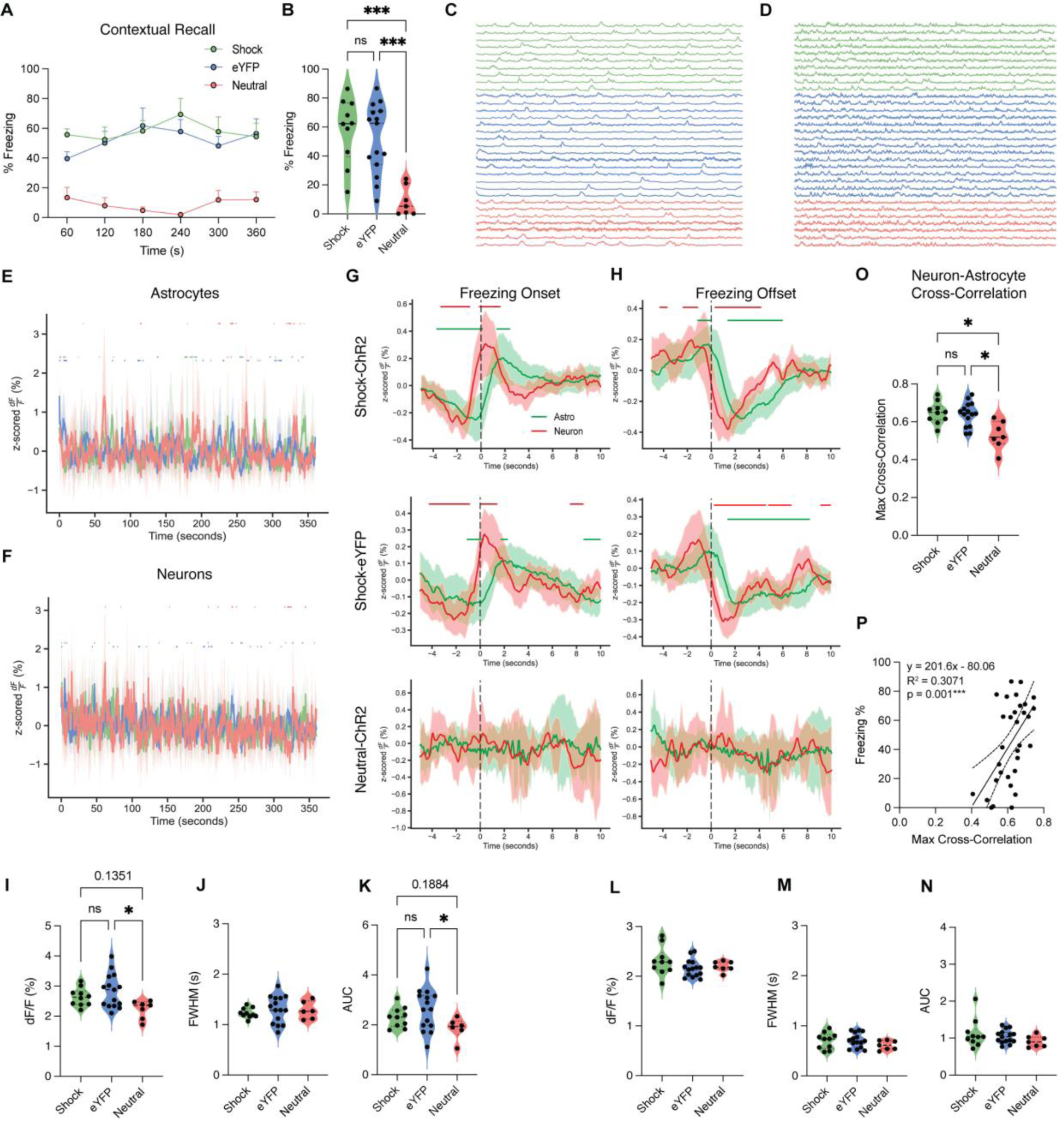
Neuronal and astrocytic calcium are responsive to freezing epochs during contextual recall in shock, but not neutral groups. (A-B) Mice in Shock-ChR2 (green) and Shock-eYFP (blue) groups expressed high levels of contextual fear recall, as shown by increased freezing percentage across the session (A) and on average (B) compared to Neutral-ChR2 (coral) mice that did not receive foot shock the day before. (C-D) Calcium timeseries for astrocytes (C) and neurons (D) in the Shock-ChR2 and Shock-eYFP groups indicated similar qualitative activity in both cell types compared to Neutral-ChR2. Each row represents a single subject across time (seconds). (E-F) Whole session peri-event analysis for astrocytic (E) and neuronal (F) calcium activity during recall showed similar activity (z-scored %dF/F) across all groups. (G-H) Peri-event analysis for neurons (red) and astrocytes (green) during freezing onset (G) and freezing offset (H) for Shock-ChR2 (top), Shock-eYFP (middle) and Neutral-ChR2 (bottom). Shock-ChR2 and Shock-eYFP mice showed a significant response across cell types to freezing onset and offset, compared to Neutral-ChR2 mice. (I-N) Calcium event metrics (z-scored) for astrocytes (I-K) and neurons (L-N); (I, L) peak height (%dF/F), (J, M) full-width half maximum (s), and (K, N) area under the curve (AUC). Astrocytes in Shock-ChR2 and Shock-eYFP groups showed significant or a trend towards an increase in peak height and AUC compared to Neutral-ChR2. Neurons showed no significant differences in any metric during recall across groups. (O) Neuron-astrocyte cross-correlation; maximum cross-correlation value (0.0-1.0) was significantly increased in both shock groups compared to neutral-ChR2 during recall. (T) Simple linear regression for average freezing percentage (%) and maximum cross-correlation value for all mice collapsed across groups during recall showed a significant relationship. For all violin plots, One-way ANOVA or Kruskal-Wallis tests were performed on normal and nonparametric data, respectively. For post-hoc multiple comparisons tests, Holm-Sidak’s (normal) or Dunn’s (nonparametric) multiple comparisons were performed with p ≤ 0.05, **p ≤ 0.01, ***p ≤ 0.001, ****p ≤ 0.0001, ns = not significant. p ≤ 0.05, **p ≤ 0.01, ***p ≤ 0.001, ****p ≤ 0.0001, ns = not significant. For event metrics and freezing behavior, shock n=10, eYFP n=15, neutral n=6-7; only one mouse in the neutral group was excluded as an outlier for astrocytic FWHM. For freezing onset and offset peri-events, shock n=10, eYFP n=15, neutral n=5; two neutral mice were excluded due to their entire lack of freezing during recall.

### Neuron-astrocyte calcium responds to freezing epochs during optogenetic reactivation of fear

To optogenetically reactivate a fear memory, mice were placed in previously neutral Cxt B for 600 seconds while receiving blue light stimulation during the 120-240 and 360-480 second time intervals. In our experiment, optogenetic reactivation drove the highest level of freezing in the Shock-ChR2 group, as expected (Figure 4A-B), though in an atypical manner from previously reported light-induced studies but nonetheless consistent with an overall increase in freezing behavior during stimulation ^32–34^. Interestingly, while we observe increases in freezing with the onset of blue light at the 120 and 360 second timepoints, the already extant freezing levels we believe reflect contextual generalization, which provides us with an opportunity to study global increases in freezing and its effects on intra-hippocampal dynamics. The Shock-eYFP group displays a moderate level of fear that is likely due to fear generalization (Figure 4A-B). As anticipated, Neutral-ChR2 mice display the lowest level of freezing, hovering around 20% on average across the session (Figure 4A-B). Qualitatively, there were no significant differences in individual nor in whole session calcium timeseries across groups for both astrocytes (Figure 4C, B) and neurons (Figure 4D-I). There were no changes in calcium event metrics across all the groups in both cell types (Figure 4F- H, J-L). Additionally, there was no significant difference in maximum cross-correlation across all the groups during the optogenetic reactivation session (Figure 4M). However, a simple linear regression revealed a significant relationship between maximum cross-correlation and average freezing during this session (Figure 4N). However, the R^2^ value for cross-correlation (R^2^ = 0.2114) was lower during this session compared to recall (R^2^ = 0.3071)(Figure 3P).

**Figure 4:**
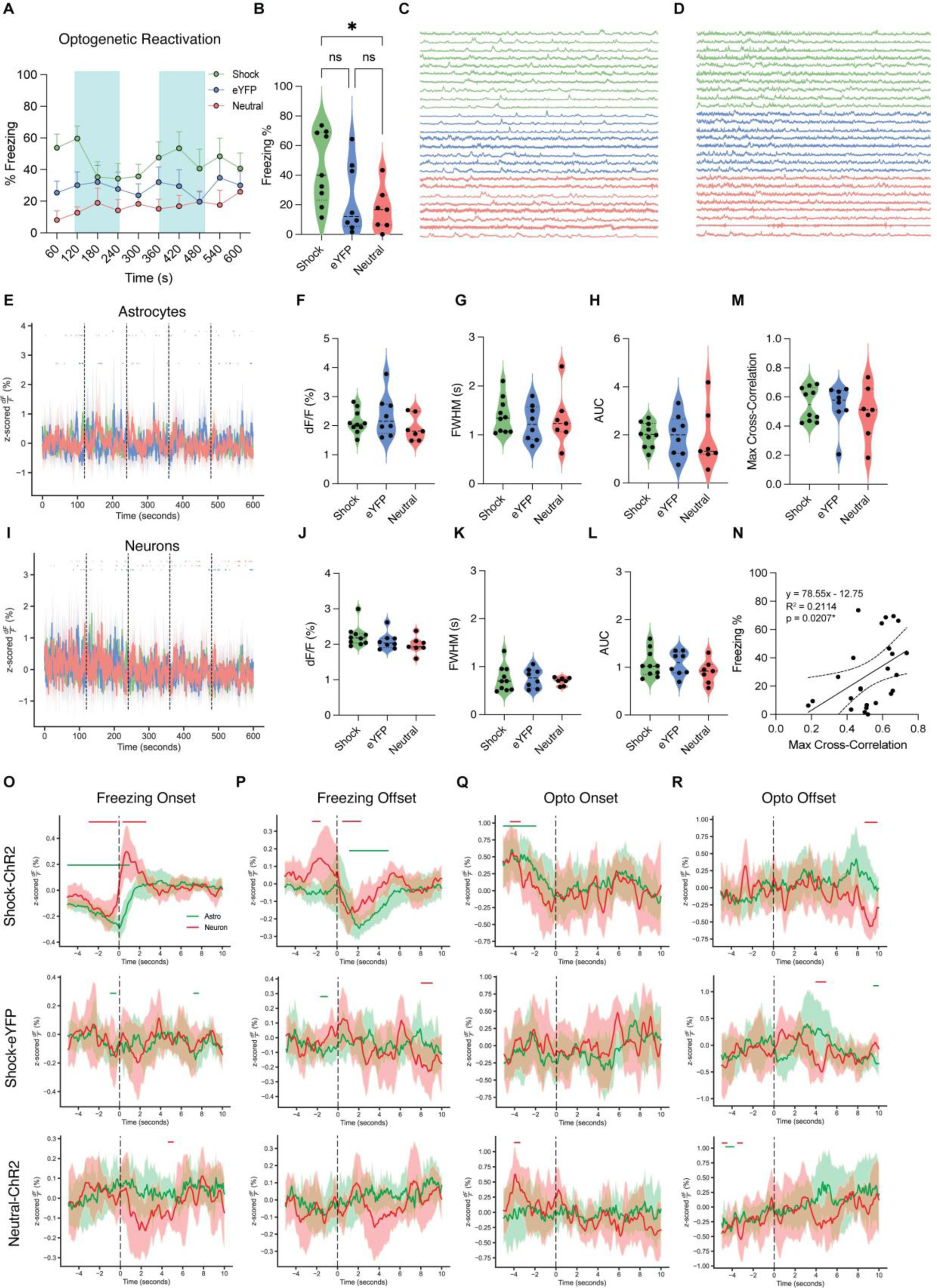
Neuron-astrocyte calcium responds to freezing epochs during optogenetic reactivation of fear. (A-B) Mice in Shock-ChR2 (green) expressed higher levels of freezing across the session (A) and on average (B) during optogenetic reactivation of a fear memory compared to Shock-eYFP (blue) and Neutral-ChR2 (coral) groups expressing moderate and low levels of fear, respectively. Blue shading indicates the light ON epochs (120-240s; 360-480s) during the 600 second session. (C-D) Calcium timeseries for astrocytes (C) and neurons (D) indicated similar qualitative activity in both cell types across all groups during optogenetic reactivation. Each row represents a single subject across time (seconds). (E, I) Whole session peri-event analysis for astrocytic (E) and neuronal (I) calcium activity during recall showed similar activity (z-scored %dF/F) across all groups. (F-H, J-L) Calcium event metrics (z-scored) for astrocytes (F-H) and neurons (J-L); (F, J) peak height (%dF/F), (G, K) full-width half maximum (seconds), and (H, L) area under the curve (AUC). Neurons and astrocytes showed no significant differences in any metric across groups. (O-R) Peri-event analysis for neurons (red) and astrocytes (green) during freezing onset (O), freezing offset (P), optogenetic stimulation onset (Q) and offset (R) for Shock-ChR2 (top), Shock-eYFP (middle) and Neutral-ChR2 (bottom). Shock-ChR2 mice showed a significant response across cell types to freezing onset and offset that is not present in other groups. (M) Neuron-astrocyte cross-correlation; maximum cross-correlation value (0.0-1.0) was not significantly different across any groups during the session. (N) Simple linear regression for average freezing percentage (%) and maximum cross-correlation value for all mice collapsed across groups during optogenetic reactivation showed a significant relationship. For all violin plots, One-way ANOVA or Kruskal-Wallis tests were performed on normal and nonparametric data, respectively. For post-hoc multiple comparisons tests, Holm-Sidak’s (normal) or Dunn’s (nonparametric) multiple comparisons were performed with p ≤ 0.05, **p ≤ 0.01, ***p ≤ 0.001, ****p ≤ 0.0001, ns = not significant. For event metrics and freezing behavior, shock-ChR2 n=10, shock-eYFP n=9, neutral-ChR2 n=7; one shock-ChR2 mouse was excluded as an outlier. For peri-events, shock-ChR2 n=10, shock-eYFP n=8, neutral-ChR2 n=7.

Interestingly, we observed coordinated calcium signatures in the Shock-ChR2 group time-locked to freezing onset and offset resembling those during natural recall in both Shock-ChR2 and Shock-eYFP mice (Figure 3G-H). Most notably, this time-locking to freezing epochs was not present in the Shock-eYFP and Neutral-ChR2 group during optogenetic reactivation (Figure 4O-P). Both the Shock-eYFP and Neutral-ChR2 exhibited some level of freezing behavior, likely due to generalization of fear, but artificial reactivation of a fear memory re-engaged astrocytes and neurons calcium activity in a stereotyped manner characteristic of natural recall. Optogenetic stimulation onset and offset did not appear to significantly change calcium activity in either cell type, although blue light onset may have led to a decrease in calcium in both cell types specific to fear memory activation (Figure 3Q-R).

### Predictive reliability of freezing is present only in the Shock-ChR2 group

As astrocytes and neurons changed their dynamics in response to freezing onset and offset during optogenetic and natural memory recall, we next if freezing could be reliably predicted from the fiber photometry traces (Figure 3G-H, Figure 4O-P). To this end, we fit a binomial generalized linear model (GLM) with a logit link to the freezing as a function of the astrocytic and neuronal photometry signals. During model selection, (*see STAR Methods*), we found that a model that used both neuronal and astrocytic signals was best able to predict freezing. Both coefficients were found to be statistically significant. We then fit this model to each animal during recall (Figure 5A-B). We found that the coefficients for the Shock-ChR2 group corresponding to the astrocytic photometry signal were not significantly different from the Shock-eYFP group but were significantly different from the Neutral-ChR2 group during natural recall (Figure 5B). Interestingly we found no significant differences between the Shock-eYFP group’s coefficients and the other groups, and we saw no significant differences in the coefficient values that correspond to the neuronal signal (Figure 5B). To assess the predictive validity of our model we used receiver operating characteristic (ROC) curves and the area under the curve (AUC) metric (Figure 5 C-D, G-H). We found that during natural recall that the Shock-ChR2 group was not significantly better at predicting freezing than the Shock-eYFP group but was significantly better at predicting freezing than the Neutral-ChR2 group (Figure 5D). Again, we found that the Shock-eYFP group was not significantly from either group. We next wanted to see how these results would change during artificial reactivation, especially as we saw a decoupling of the peri-event responses to freezing onset and offset between the Shock-eYFP groups during artificial reactivation (Figure 4O-P). Furthermore, we found that the astrocyte signal corresponding beta coefficients were significantly different in the Shock-ChR2 group compared to both other groups (Figure 5F). We again observed no statistical differences in the neuronal signal coefficients (Figure 5F), and then found that in the AUC metrics the Shock-ChR2 group had significantly better predictive power than both the Shock-eYFP and Neutral-ChR2 groups (Figure 5H). Overall, this analysis suggests that our optogenetic stimulation paradigm preserves freezing-specific information within the photometry signals of vCA1 neurons and astrocytes.

**Figure 5.**
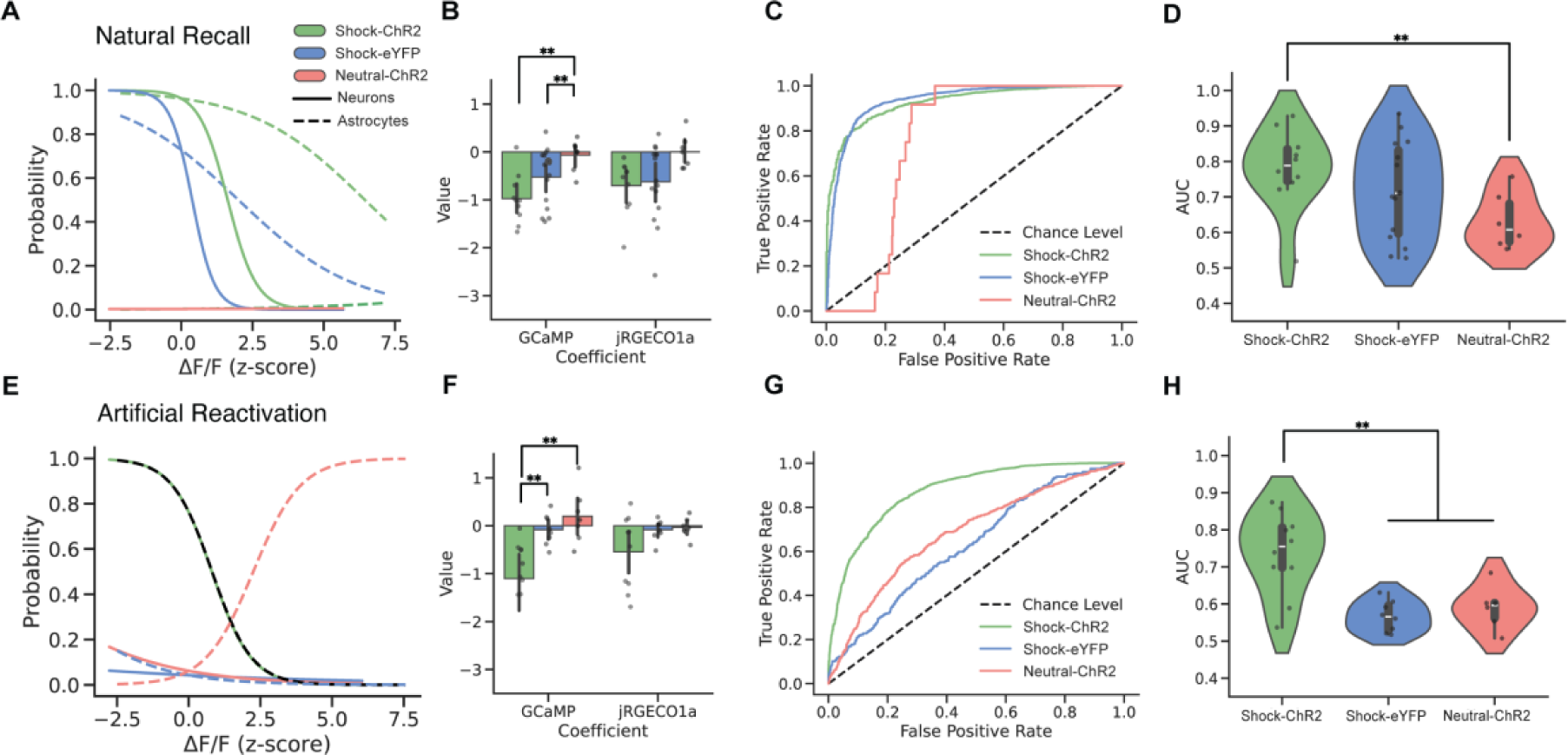
Optogenetic reactivation of a hippocampal engram preserves cellular signals of fear. (A) Best-fitting model for each group during natural recall. Lines reflect partial effects where the other signal is held at the mean value to show how the probability of freezing changes as a function of signal value. (B) Values of beta coefficients for astrocyte and neuronal photometry signals in the regression models during natural recall. (C) Receiver operating characteristic (ROC) plot for the best fitting model for each group. (D) Area under the curve (AUC) values for each model across groups (E) Best-fitting model for each group during artificial reactivation. Lines reflect the partial effects where the other signal is fixed to the mean. (F) Beta coefficients for each signal across groups during artificial reactivation. (G) ROC plot for the best fitting model for each group. (H) AUC values for each model across groups.

### Mice display low freezing and no differences in calcium dynamics across all groups during habituation

On Day 1, mice were placed in a neutral Cxt B for a habituation session while receiving blue light stimulation during the 120-240 and 360-480 second time intervals. As expected, mice exhibited low levels of freezing because no cells were tagged with ChR2 (Supplemental Figure 1A-B). Calcium timeseries and whole session peri-event analysis for astrocytes (Supplemental Figure 1C-E) and neurons (Figure 3D, I) showed qualitatively similar activity in all groups during habituation. Calcium event characteristics were calculated for all groups for both neurons and astrocytes. In both cell types, there were no significant differences in any event metrics: peak height, area under the curve, and full-width half-maximum (Supplemental Figure 1F-M, J-L). Further, we calculated neuron-astrocyte cross-correlation, observing, as expected, non-significant differences across the three groups (Supplemental Figure 1M). We fit a simple linear regression to test if differences in cross-correlation can predict the animal’s average freezing level. During the habituation session, we did not observe a significant relationship between cross-correlation and freezing (Supplemental Figure 1N). Next, we performed peri-event analysis to examine astrocytic and neuronal calcium dynamics at the timepoints of freezing initiation and termination, and the onset and offset of blue-light stimulation. We do not observe the same stereotyped decreases and increases in calcium time-locked to freezing bouts observed during natural or artificial fear memory recall (Supplemental Figure O-P). This confirms that the calcium signatures we observe during recall for both shock groups and artificial fear memory recall in the Shock-ChR2 group are unique to fear memory engagement. Blue-light stimulation onset and offset during habituation did not reliably time-lock to calcium in any group, with significant events likely related to spurious, noisy activity that we commonly observe during ‘neutral’ contexts (Supplemental Figure Q-R) ^31^.

## Discussion

### The role of neuron-astrocyte dynamics during natural and artificial fear memory

In this study, we asked how artificial reactivation of fear ensembles affects intrahippocampal neuronal and astrocytic calcium activity and how this interfaces with fear behavioral states. Our data demonstrates the emergence of coordinated calcium signatures time-locked to onset and offset of freezing during CFC that become more distinctive during natural recall in the Shock-ChR2 and -eYFP groups. This is consistent with previous findings from our lab showing that astrocytes in the basolateral amygdala are time-locked to freezing epochs during CFC and recall. Optogenetic reactivation of a dDG fear engram recapitulates these dynamics during freezing bouts, whereas stimulation of neutral-tagged neurons (Neutral-ChR2) and control Shock-eYFP mice do not, suggesting that this manipulation mimics the natural expression of fear.

### Optogenetic stimulation of a fear engram elicits atypical behavioral response

Based on previous work and the work of others, when mice with tagged Shock-ChR2+ cells undergo optogenetic stimulation of these fear-related neuronal ensembles -- they display light-induced freezing behavior. For this reason, we expected that our mice in the Shock-ChR2 group would display high levels of freezing during the light ON epochs and decrease during light OFF epochs. This ‘see-saw’ effect is not typically observed in Shock-eYFP and Neutral-ChR2 groups where they display lower levels of freezing overall. This is because the eYFP+ cells are not blue-light sensitive, and thus should not be ‘reactivated’ or brought back online to drive fear, and the Neutral-ChR2 group has a non-shock-related experience labeled. Our behavioral findings during optogenetic reactivation showed an increase in freezing with the onset of blue-light stimulation that was modest in the Shock-ChR2 group. In line with our observations, previous engram research has shown a range of variability in light-induced freezing behaviors driven by differences in stimulation frequency and brain region ^35^. For instance, recent work has shown a very modest increase between light-ON and OFF epochs (e.g. ∼ 5% changes in freezing) with optogenetic stimulation of the prefrontal cortex (PFC), which is consistent with our findings ^32^. This is further supported by other work showing large variability in freezing, with the distributions between shock and no shock largely overlapping and few mice driving this significant difference during the light-ON periods ^33^. Further, when collapsed across the entire optogenetic session (light-ON and light-OFF epochs together for each group), this study observed no significant differences across shock and no shock mice. Both of these works nonetheless successfully dissociated light-induced changes in behavior from light-induced changes in cellular dynamics. Importantly, our optogenetic stimulation, although similarly modest in behavioral effects, indeed recapitulates coordinated calcium signatures resembling natural recall that are time-locked to freezing behavior, only in the Shock-ChR2 group. This may suggest that during fear states, and specifically during bouts of freezing, fear engram reactivation alters downstream neuronal and astrocytic calcium dynamics and this alteration mimics natural fear states.

### Correlation between neuronal and astrocytic calcium can predict fear states

Due to the robust coordinated responses between neurons and astrocytes at the onset and offset of freezing epochs, we next tested if the two cell types become more correlated with an increase in average fear. Our analysis showed no significant differences in maximum cross-correlations between groups during CFC. This is likely due to the emergence of the fear state across the session and may require consolidation of the memory ^32^. In line with this hypothesis, in natural recall there was increased correlation between neurons and astrocytes in the shocked groups compared to the no-shock group. A simple linear regression analysis revealed that cross-correlations had a linear relationship to average freezing in the session. During the artificial recall session, although the Shock-ChR2 group exhibited increased freezing levels and stereotyped calcium signatures time-locked to freezing characteristic of natural fear, however, there were no significant differences in maximum cross-correlations. Moreover, we observed more variability in cross-correlations between animals in both shock groups compared to natural recall. We speculate that optogenetic stimulation of an engram multiple synapses upstream^2^ alters natural dynamics and leads to decreased coordination between cell types across the session compared to recall. Despite the induction of elevated fear levels and stereotyped calcium time-locking to freezing epochs during optogenetic reactivation, this artificial perturbation is likely to produce more variability, both behaviorally and cellularly.

### Statistical Modelling Enhances Interpretability of Engram Research

The ability to manipulate engrams via inhibition or activation has been technologically feasible for the past two decades ^5,36^, but the majority of research relies on using behavioral metrics (e.g. freezing) and or measurements of IEG expression as the primary readouts. However, as real-time recording and concurrent optogenetic perturbations are becoming more prevalent, a higher level of cellular and behavior resolution is warranted to measure phenotypes in an unbiased manner. One caveat with using a single behavioral dimension to assay memory is that it makes a *prima facie* assumption that subjects are independent and identically distributed. However, several studies have shown that metrics including freezing are not always universally expressed and show considerable variability. For instance, the often-observed “bi-directional freezing” often observed during optogenetic stimulation is not always present ^32–34^. In this study, we propose that as real-time recording approaches become more prevalent in engram research, statistical modeling will become indispensable. We used a classic statistical model, the Generalized Linear Model, which has been seminal in many other fields of neuroscience ^37^. We found that using our two photometry signals from astrocytes and neurons reliably predicted freezing behavior in mice, even though we observed non-traditional freezing curves. Accordingly, we speculate that engram stimulation preserves encoded “fear” states within the activity of vCA1 neurons and astrocytes. This suggests that while engram stimulation is undoubtedly an artificial approach, we nonetheless observe physiological responses expected under natural fear memory recall. We believe that this result paired with our observation of significant peri-events in the Shock-ChR2 group only during optogenetic stimulation advances the idea that our stimulation is recapitulating fear-related neuronal responses. It is worth underscoring that our model selection procedure suggests that neuronal signals are playing an important predictive role in our model, despite the absence of significant group level differences in the corresponding beta coefficients. We believe there are a few reasonable explanations that could resolve this conflict. For example, like other red calcium indicators, jRGECO1a has lower signal-to-noise ratio than its GCaMP6f counterpart, and so it is feasible that some animals will have much more reliable predictive capabilities in this channel than others. An additional caveat is that we fit a separate model to each of the animals and assume each mouse is independent. Thus, future research could augment our approach in at least two manners: one could use a generalized linear-mixed effects model which would supply a maximum a posteriori estimate of the posterior distribution of the beta-coefficients; or, one could use a fully Bayesian approach and partial-pool all of the data to take into account the inherent variability of different mice. Regardless, we believe that even simple statistical models, such as the ones used here, will enhance the interpretation of future engram studies.

### Advancing the study of natural and artificially-driven memories

Our study demonstrates that optogenetic reactivation of a fear engram induces neuronal-astrocytic dynamics that resemble the cellular responses observed during natural fear memory recall. Interestingly, recent work has characterized the relationship between phase-specific optogenetic stimulation of the DG and theta oscillations in the dorsal CA1 of the hippocampus using local-field potentials. Here, they find that stimulation during the trough of theta is more effective at driving freezing behavior compared to the peak of theta. The variability in freezing behavior in our data could be potentially explained by imprecise stimulation patterns that do not yet take phase-specific relationships between engram activation and ongoing rhythms in the brain into account ^34^. This study also did not observe significant light-induced freezing during light-ON epochs at 20Hz in the dDG, but did observe this during trough stimulation, further supporting the notion that the brain contains optimal rhythmic windows during which optogenetic stimulation of an engram may be most effective at driving behavior. This work builds on others that have recorded from within or downstream of DG engram stimulation. For example, calcium recordings of *F*-RAM and *N*-RAM ensembles expressing GCaMP after CFC revealed that they were differentially reactivated during recall in the DG, with the *N*-RAM ensemble uniquely engaging for memory discrimination^38^. Further, single-unit electrophysiological activity in the BLA of mice while simultaneously activating DG positive engram cells had both excitatory and inhibitory effects on individual cells^7^. Finally, 20 Hz stimulation of a sparse population of DG granule cells was shown to excite and inhibit an equal number of neurons downstream in CA3, which provides a putative mechanism for how both excitation and inhibition work together to effectively express a memory^39^. Together with this previous work, our study supports the idea that optogenetic reactivation is sufficient to induce behavioral and cellular fear states similar to those observed during natural recall.

## Author Contributions

Conceptualization, R.L.S, R.A.S, S.R.; Methodology, R.L.S, R.A.S, S.R.; Formal Analysis, R.L.S, R.A.S, M.D.B; Investigation, R.L.S, R.A.S, M.D.B, A.D., A.Y.P.; Visualization, R.L.S, R.A.S; Writing – Original Draft, R.L.S, R.A.S, M.D.B, S.R.; Writing – Review & Editing, R.L.S, R.A.S, M.D.B, S.R.; Funding Acquisition, S.R.; Resources, S.R. Supervision, S.R.

## Acknowledgements

We thank members of the Ramirez laboratory for their helpful comments and suggestions on the manuscript. This material is based upon work supported by the Air Force Office of Scientific Research (AFOSR) under award number FA9550-21-1-0310, the Ludwig Family Foundation, an NIH Early Independence Award DP5 OD023106-01, NIH Transformative Award, Brain and Behavior Research Foundation Young Investigator Grant, McKnight Foundation Memory and Cognitive Disorders Award, Pew Scholars Program in the Biomedical Sciences, the Chan-Zuckerberg Initiative, and the Center for Systems Neuroscience and Neurophotonics Center at Boston University. Behavioral schematics were created using BioRender.com

## Main Figure Titles & Legends

**Supplemental Figure 1 (related to Figure 4):**
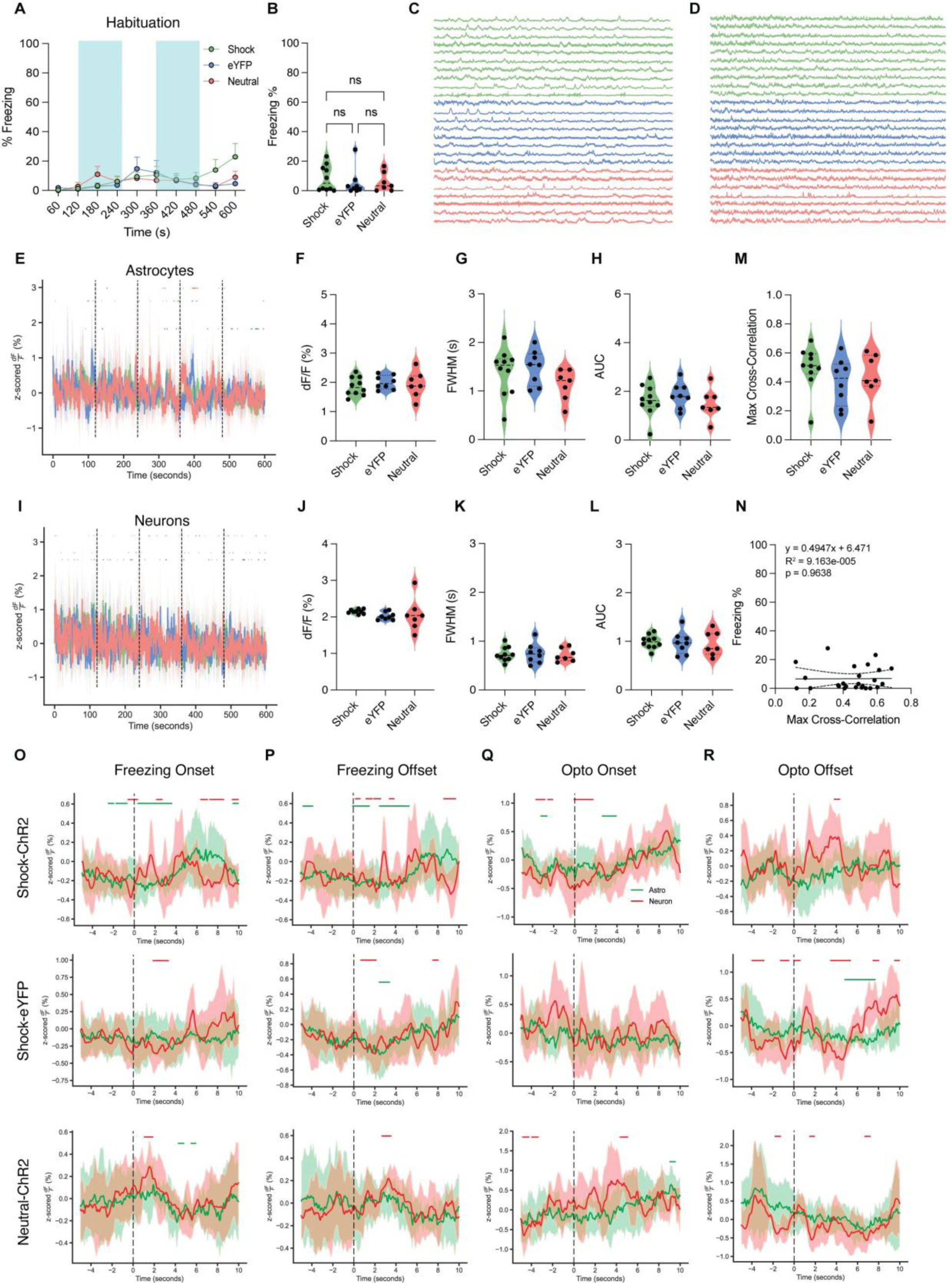
Mice display low freezing and no differences in calcium dynamics across all groups during habituation. (A-B) Mice in all groups, Shock-ChR2 (green), Shock-eYFP (blue) and Neutral-ChR2 (coral) expressed low levels of fear, as evidenced by freezing percentage across the session (A) and on average (B). Blue shading indicates the light ON epochs (120-240s; 360-480s) during the 600 second session. (C-D) Calcium timeseries for astrocytes (C) and neurons (D) indicated similar qualitative activity in both cell types across all groups during habituation. Each row represents a single subject across time (seconds). (E, I) Whole session peri-event analysis for astrocytic (E) and neuronal (I) calcium activity during habituation showed similar activity (z-scored %dF/F) across all groups. (F-H, J-L) Calcium event metrics (z-scored) for astrocytes (F-H) and neurons (J-L); (F, J) peak height (%dF/F), (G, K) full-width half maximum (s), and (H, L) area under the curve (AUC). Neurons and astrocytes showed no significant differences in any metric during the session across groups. (O-R) Peri-event analysis for neurons (red) and astrocytes (green) during freezing onset (O), freezing offset (P), optogenetic stimulation onset (Q) and offset (R) for Shock-ChR2 (top), Shock-eYFP (middle) and Neutral-ChR2 (bottom). All groups showed significant periods of time for either cell type, but no evident pattern of activation occurred during each event. (M) Neuron-astrocyte cross correlation; maximum cross-correlation value (0.0-1.0) was not significantly different across any groups during the session. (N) Simple linear regression for average freezing percentage (%) and maximum cross-correlation value for all mice collapsed across groups during habituation showed no significant relationship. For all violin plots, One-way ANOVA or Kruskal-Wallis tests were performed on normal and nonparametric data, respectively. For post-hoc multiple comparisons tests, Holm-Sidak’s (normal) or Dunn’s (nonparametric) multiple comparisons were performed with p ≤ 0.05, **p ≤ 0.01, ***p ≤ 0.001, ****p ≤ 0.0001, ns = not significant. For event metrics and freezing behavior, shock-ChR2 n=10, shock-eYFP n=8, neutral-ChR2 n=7; one shock-ChR2 mouse was excluded as an outlier for neuron peak height. For peri-events, shock-ChR2 n=10, shock-eYFP n=8, neutral-ChR2 n=5; two neutral-ChR2 mice were removed due to a lack of freezing epochs.

## STAR METHODS

### Resource Availability

Further information and requests for resources and reagents should be directed to and will be fulfilled by the lead contact, Steve Ramirez.

### Materials Availability

This study did not generate new unique reagents.

### Data and Code Availability

All data reported in this paper will be shared by the lead contact upon request. All original code has been deposited at https://github.com/rsenne/RamiPho and is publicly available as of the date of publication.

### Experimental model and subject details

#### Mice

Wild type, male C57BL/6J mice (P29-35; weight 17-19g; Charles River Laboratories strain #027) were housed in groups of 3-5 mice per cage. The animal facilities (vivarium and behavioral testing rooms) were maintained on a 12:12 hour light cycle (0700-1900). Mice received food and water *ad libitum* before and after surgery. All mice were placed on a 40mg/kg doxycycline (Dox; Bio-Serv) diet 48 hours prior to surgery to inhibit any ‘tagging’ that could occur with the infusion of the Tet-tag viral cocktail. Following surgery, mice were group-housed with littermates and allowed to recover for 4 weeks before experimentation for expression of genetically-encoded calcium indicators (GECIs). All subjects were treated in accord with protocol 201800579 approved by the Institutional Animal Care and Use Committee (IACUC) at Boston University. As a limitation of this study, only male mice were utilized, and future work will investigate if these findings generalize to females as well.

#### Stereotaxic Surgery

For all surgeries, mice were initially anesthetized with 3.0-3.5% isoflurane inhalation during induction and maintained at 1-2% isoflurane inhalation through stereotaxic (David Kopf Instruments) nose cone delivery (oxygen 1L/min). Ophthalmic ointment was applied to the eyes to provide adequate lubrication and prevent corneal desiccation. The hair on the scalp above the surgical site was removed using Nair hair removal cream and subsequently cleaned with alternating applications of betadine solution and 70% ethanol. 2.0% lidocaine hydrochloride (HCl) was injected subcutaneously as local analgesia prior to midsagittal incision of the scalp skin to expose the skull. 0.1mg/kg (5mg/kg) subcutaneous (SQ) dose of meloxicam was administered at the beginning of surgery. All animals received craniotomies with a 0.5-0.6 mm drill-bit for ventral CA1 (vCA1) and dorsal dentate gyrus (dDG) injections and implants.

A 10μL airtight syringe (Hamilton Company) with an attached 33-gauge beveled needle was slowly lowered to the coordinates of hippocampal ventral CA1 (vCA1): −3.16 anteroposterior (AP), −3.10 mediolateral (ML) and −4.50 dorsoventral (DV) for fiber photometry recordings. All coordinates are given relative to bregma (mm). A volume of undiluted 250nL:500nL AAV5-GfaABC1D-cyto-GCaMP6f-SV40 (AddGene #52925)^40^ and AAV9-hSyn-NES-jRGECO1a-WPRE.SV40 (AddGene #100854)^41^ was injected using a microinfusion pump for the vCA1 coordinate at 50nL/min (UMP3; World Precision Instruments). After the injection was complete, the needle remained at the target site for 7-10 minutes post-injection before removal. Following viral injection, a unilateral optic fiber (200μm core diameter; 1.25mm ferrule diameter, NA=0,.37, length = 4.5mm; Neurophotometrics) was implanted at −4.60DV, slightly below the site of viral injection.

To enable engram tagging and manipulation via optogenetics, bilateral dorsal dentate gyrus (dDG) was infused with 1:1 undiluted AAV9-c-fos-tTA-BGHpa and AAV-TRE-ChR2-eYFP/AAV9-TRE-eYFP (UMass Vector Core, custom) to label neuronal ensembles with channelrhodopsin (ChR2), a blue-light sensitive protein. A volume of 250nL of the viral cocktail was infused at 100nL/min into bilateral dDG at −2.20 AP, 土 1.30 ML and −2.0DV. Mice received bilateral optic fiber implants 0.2mm above the site of infusion (1.80 DV)(Doric Lenses). The implant was secured to the skull with a layer of adhesive cement (C&M Metabond) followed by multiple layers of dental cement (Stoelting). Following surgery, mice were injected with a 0.1mg/kg intraperitoneal (IP) dose of buprenorphine. They were placed in a recovery cage with a heating pad until fully recovered from anesthesia. Histological assessment verified viral targeting and data from off-target injections were not included in analyses.

#### Tet-Tag System

The Tet-tag system is an inducible, activity-dependent labeling strategy that relies on the neuronal expression of the immediate-early gene, *cfos*. This system is composed of a viral cocktail of c-fos-tTA and TRE-ChR2-eYFP or eYFP control fluorophore. This genetic strategy has been used to label (‘tag’) a neuronal ensemble, typically referred to as a memory *engram*, that contains information that is vital to the encoding and recall of a recent experience. This system couples the *cfos* promoter to the tetracycline transactivator (tTA). In its protein form, tTA directly binds to the tetracycline response element (TRE) in a doxycycline (Dox)-dependent manner and drives expression of a protein of interest (i.e. channelrhodopsin (ChR2) and/or fluorophore). This allows for the temporal regulation of a ‘tagging’ window when removed from an animal’s diet 48 hours prior to a salient experience (i.e. contextual fear conditioning or home cage exposure). Returning the mice to Dox diet immediately after the experience of interest ‘closes’ the tagging window, and they remain untouched for 24 hours to prevent off-target labeling. Most importantly, the expression of ChR2 allows for optogenetic reactivation of the experience tagged during encoding. Specifically, optogenetic stimulation of ChR2+ negative engram cells with blue light in a ‘neutral’ context manifests as a freezing response, typically referred to as *light-induced freezing*.

#### Fiber Photometry Data Collection

A 470-nm LED (Neurophotometrics; FP3002) delivered an excitation wavelength of light to astrocytes expressing GCaMP6f via a single fiber optic implant. The emitted 530-nm signal from the indicator was collected via this same fiber and patch cord (Doric Lenses), spectrally-separated using a dichroic mirror, passed through a series of filters, and was focused on a scientific camera. Isosbestic signals were simultaneously captured by alternating excitation with 415-nm LED to dissociate motion, tissue autofluorescence, and photobleaching from true changes in fluorescence. For these dual-color experiments, we acquired data simultaneously from two channels by adding a 560-nm LED to excite jRGECO1a. All wavelengths were interleaved and collected simultaneously using Bonsai interfacing with the Neurophotometrics system ^42^. The sampling rate for the calcium signals was 10 Hz per channel.

#### Behavioral Testing

On Day 1, mice were habituated for 600s to a neutral context (Cxt B) while undergoing optogenetic stimulation to control for encoding of a novel environment and light-stimulation alone (Coulbourn Instruments). Because they are still consuming a Dox diet, there should be no experience or engram ‘tagged’ during this time. After this session, the Dox-containing diet was replaced with standard mouse chow *(ad libitum*) 48 hours prior to behavioral tagging to open a time window of activity-dependent labeling. On Day 3, mice were placed into the shock context (Cxt A) where they underwent CFC for 360s. Foot shocks (0.75mA, 2s duration) were administered at the 120s, 180s, 240s and 300s time points and animals were immediately placed back on Dox diet for 24 hours, closing the tagging window. This labeled sufficiently active cells with ChR2-eYFP or eYFP alone. On Day 4, mice were placed back in Cxt A where they received foot shocks on the previous day for 360s of ‘natural recall’. On the following Day 5, ChR2- eYFP+ or eYFP+ cells were optogenetically stimulated (450nm laser diode, 20Hz, 10ms pulse, 15mW output) in Cxt B with alternating 2-minute light-ON and light-OFF epochs [off/on/off/on/off] for a total of 600s (Doric Lenses). 90 minutes after the start of the last behavioral session, we performed perfusions to measure endogenous c-Fos at its peak, providing a proxy of recent neural activity resulting from optogenetic reactivation of dDG. Brains were sliced on a vibratome at 50 um thickness, immunohistochemistry, and confocal microscopy (Zeiss LSM800, Germany) were performed to quantitatively analyze the total number of ‘tagged’ cells in dDG, number of overlaps between endogenous c-Fos+ and ‘tagged’ ChR2/eYFP+ cells and expression profile of GCaMP6f and jRGECO1a. Time series data were analyzed in vCA1 from neurons and astrocytes across all experimental days.

All of these sessions took place in mouse conditioning chambers (Coulbourn Instruments) with metal-panel side walls, plexiglass front and rear walls and a stainless-steel grid floor composed of 16 grid bars (18.5 x 18 x 21.5cm). The grid floor was connected to a precision animal shocker to deliver 2 second duration, 0.75mA foot shocks. Context A was composed of this standard fear conditioning chamber with white light and no odorants. Context B was in a separate room, with a textured black floor to cover the shock grid, striped laminated walls, an orange odor, and red light in the front of the room. No auditory changes were made across contexts. A web camera was mounted in front of the chamber to record animal behavior that was triggered by the onset of calcium recording in Bonsai/Neurophotometrics. The behavioral session was triggered by a computer running FreezeFrame4 software (Actimetrics). The chambers were cleaned with 70% ethanol solution prior to each animal placement.

#### Immunohistochemistry

On the final day of behavior, mice were overdosed with 3% isoflurane and perfused transcardially with cold (4°C) 1 X Phosphate-buffered saline (PBS; Gibco) followed by 4% Paraformaldehyde (PFA; pH = 7.4; Sigma-Aldrich) in PBS. Brains were extracted and kept in PFA at 4°C for 24-48 hours. Brains were sectioned into 50μm thick coronal sections with a vibratome and collected in cold PBS or 0.01% sodium azide in PBS for long-term storage. Sections were washed three times for 10-15 minutes with PBS or PBST to remove 0.01% sodium azide used for storage. Vibratome sections were incubated for 2 hours in PBS combined with 0.2% Triton (PBST; Teknova) and 5% bovine serum albumin (BSA; Sigma-Aldrich) on a shaker at room temperature for blocking. Sections were incubated in the primary antibodies (1:1000 rabbit polyclonal anti-cFos [SySy; #226 008]; 1:1000 chicken polyclonal anti-GFP [Invitrogen; #A10262]; 1:1000 guinea pig anti-RFP [SySy; #390 004] diluted in PBS/1% BSA/Triton X-100 solution at 4°C for 24- 48 hours depending on the stain of interest. The slices were washed three times for 10-15 minutes each in 1xPBS or 0.2% PBST. The secondary antibodies were diluted in secondary antibody solution (PBS/1% BSA/Triton X-100) and incubated for 2 hours at room temperature. The following secondary antibodies were used: 1:200 Alexa goat anti-rabbit 555 [Invitrogen; #A-21428]; 1:200 Alexa goat anti-chicken 488 [Invitrogen; #A-11039]; 1:500 Alexa goat anti-guinea pig 555 [Invitrogen; #A-11073]. The sections were then washed three times with 1xPBS or PBST for 10-15 minutes each and mounted using Vectashield HardSet Mounting Medium with DAPI (Vector Laboratories Inc.). Once dry, slides were sealed with clear nail polish on each edge and stored in a slide box in the fridge (4°C). Mounted slices were imaged using a confocal microscope (Zeiss LSM800, Germany). Brains from all mice used in fiber photometry experiments were analyzed to check adequate fiber location and proper and selective viral expression. Animals that did not meet the criteria for proper fiber location and virus expression were discarded.

#### Image Acquisition

All coronal brain slices were imaged through a Zeiss LSM 800 epifluorescence microscope with a 20x/0.8 numerical aperture objective using Zen2.3 software. Brains from all mice used in fiber photometry experiments were analyzed to check adequate fiber location and proper and selective viral expression. Animals that did not meet the criteria for proper fiber location and virus expression were discarded.

Images of the dDG were captured in a 2 x 4 tile (1280 x 640 um) z-stack. DAPI, cFos and green fluorescent protein (GFP) were imaged as separate channels for target verification, ensemble size quantification and overlaps of ‘reactivation.’ 3-4 slices (6-8 dDG ROIs) were imaged for each animal for averaging.

#### Quantification and Statistical Analysis

All details of statistical analysis (statistical test used, n value, comparisons, test statistics, p-values, post-hoc multiple comparisons, outliers removed, and results of normality and variance measures) can be found in the Supplemental Statistical Table. Brief notes of statistical tests are included in the main and supplemental figure legends (statistical test, n value, outliers or mice removed).

##### Cell Counting Analysis

dDG images were processed prior to quantification in FIJI (ImageJ). Images were processed using a custom macro to split channels, adjust brightness/contrast and z-project (maximum intensity). Regions of interest were selected using the Polygon tool so that only cells within the dDG granule cell layer were quantified. cFos, GFP and DAPI channels were separately quantified in Ilastik ^43^, a supervised machine-learning analysis tool. This method uses a pixel and object classification pipeline facilitated by a human annotator and allows for automated batch processing once the algorithm has been properly trained. Overlaps between cFos and GFP were quantified manually in FIJI (ImageJ) using the Cell Counter plug-in due to the small number of cells.

Once cells were counted, the amount of cFos+, GFP+ and cFos+GFP+ (overlap) cells in each slice were normalized to the total number of DAPI+ cells (cFos+/DAPI+, GFP+/DAPI+ and overlap/DAPI). The chance of an overlap was defined as (cFos+/DAPI+) x (GFP+/DAPI+), which was calculated for each slice. Overlap/chance was then calculated by dividing overlap/DAPI by chance for each individual slice. Overlap/chance was averaged across all slices for each mouse to generate a single average value that was used in statistical analysis.

##### Fiber Photometry Analysis

All fiber photometry analysis was performed using an in-house pipeline available at https://github.com/rsenne/RamiPho. Extracted photometry signals first underwent baseline correction using the adaptive iteratively reweighted Penalized Least Squares (airPLS) algorithm. For this algorithm we set *λ* = 10.7 and *p* = 0.05 with a maximum of 100 iterations. Following baseline correction, we used a Kalman Filter to smooth each trace ^44^. The Kalman Filter (or Gaussian Linear Dynamical System) is a model of the form:

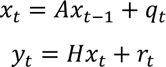

Where *A* is the state transition matrix, *H* is the observation model matrix, *q*_*t*_ is the process noise which is zero-mean Gaussian distributed with *Q*_*t*_ covariance, *Q*_*t*_: *q*_*t*_ ∼ *N*(0, *Q*_*t*_), and *r*_*t*_ is the observation noise which is zero-mean Gaussian noise with covariance *R*_*t*_: *r*_*t*_ ∼ *N*(0, *R*_*t*_). To use the Kalman Filter we decided to use a p-order Autoregressive (*AR*(*p*)) process as a process model. We found that almost all photometry traces were well described by an *AR*(3) model by plotting the partial autocorrelation functions and determining how many lags were outside of the theoretical 95% rejection region. To this end, we modeled every photometry trace as an *AR*(3) process of the form:

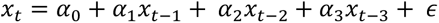

Where *x*_*t*_ is the current time step, *x*_*t*−*k*_ is the k-th previous timestep, *α*_0_ − *α*_3_ are constant coefficients and epsilon is a Gaussian distributed error term. Thus, for our state transition matrix and observation model matrix we used:

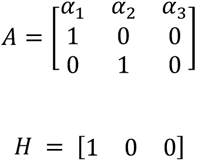

For the model we set the initial covariance as the identity matrix *I* (3) and the initial state as the zero vector (0, 0, 0). We used the expectation-maximization algorithm to find the initial process noise and observation noise covariance matrices. We then used the Kalman (Rauch–Tung–Striebel) Smoother and then used the smoothed estimate of the hidden continuous state as our photometry trace. After smoothing, we then calculated 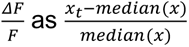 and also a z-scored version as 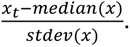 Following this calculator, we then fit an ordinary least squares regression model to correct for motion and artifact noise in the recording according to the following model:

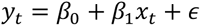

Where y_t is the calcium indicator x_t is the isobestic channel the \beta are constant coefficients and epsilon is Gaussian distributed noise. After fitting this model, we used the residuals as our motion corrected trace.

For event detection we used the find_peaks() function in Scipy. Two researchers manually choose parameters across all traces until they were both in agreement. Importantly, all parameters were chosen before any statistical or exploratory analysis was performed and were not altered at a later time point.

For event-triggered average significance, we used a tCI confidence interval method previously proposed^45^. For our event triggered averages we aggregated all within-animal signals so that we could assume that our samples were independent and identically distributed. Thus, for this method we assumed each time point was distributed according to a student-t’s distribution. We then marked any period of time greater than 0.8 seconds (this decision was arbitrary and could be chosen to be longer for more conservative estimation, but this is longer than the proposed time threshold in the original paper) that did not include the baseline of 0 (traces were median-shifted to zero) was marked at a significant peri-event.

To assess how neural and astrocytic traces could be used to predict freezing we used a Binomial GLM with the canonical logit link function. For model selection, all data was pooled together on the Shock-ChR2 animals, and we performed 10-fold cross validation and used the Maximum Likelihood Ratio test to assess the following models 1) just a constant, 2) a constant and the astrocyte signal, 3) a constant and the neuronal signal, 4) and a constant, the astrocyte signal, and the neural signal. The MLRT results suggested that the full model was superior to all of the nested models. Our cross-validation approach also found that the full model was the most generalizable. After this model selection routine, we fit the chosen model to each animal’s data separately.

##### Behavioral Analysis

To perform unbiased behavioral evaluation, the pose estimation algorithm, DeepLabCut, was used for animal kinematics (position, acceleration, velocity)^46^. This open-source toolbox allows for training of a deep neural network using a small number of behavioral videos. This method was confirmed for accuracy by a blinded researcher that manually scored a subset of videos. Additionally, Any-Maze (Stoelting Co.) was used for supervised automated analysis of freezing bout initiation and termination. This behavioral data was time locked to our fiber photometry time series data for analysis.

To extract acceleration and velocity information from DeepLabCut, we again used the Kalman Filter as described above. We first calculated the center of mass of the pose estimates for the mice to estimate the x and y coordinates. After this we then formulated a Kalman Filtering model using the following matrices:

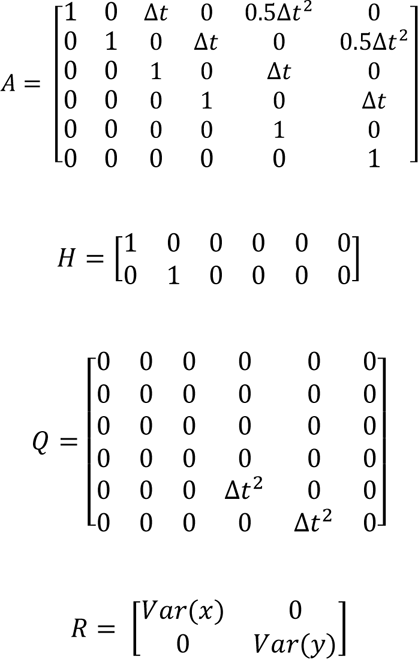

For the initial state mean we used: (*x*_1_, *y*_1_, 0, 0, 0, 0) and for the initial covariance we used the identity matrix: *I*(6). Using this method, we were able to extract smooth estimates of the (x, y) position, velocity, and acceleration.

##### Statistical Methods

Statistical analyses were performed using Python and GraphPad Prism v10. For behavioral measures and event metrics, violin plots show the data distribution with dashed lines indicating the upper and lower quartiles and the solid line indicating the median. All data were checked for normality using Shapiro-Wilks and Kolmogorov-Smirnov tests, equality of variance (standard deviation) using Brown-Forsythe test and outliers using the ROUT method. This method was recommended by GraphPad Prism, which uses identification from nonlinear regression. We chose a ROUT coefficient Q value of 10% (False Discovery Rate), making the threshold for outliers less-strict and allowing for an increase in power for outlier detection. Mice that were excluded for other reasons, such as a lack of freezing epochs, were noted in the figure legends where applicable in peri-event analysis.

To analyze differences between groups for behavioral measures, event metrics and cross-correlation measures, we used: One-way ANOVAs (normal) or Kruskall-Wallis test (nonparametric) with Holm-Sidak’s (normal) or Dunn’s (nonparametric) post-hoc multiple comparisons tests, if applicable. If variances were not equal, a Brown-Forsythe ANOVA (normal) was performed instead with Dunnett’s T3 multiple comparisons test, if applicable. To analyze differences between groups and across time within a single session we used: Two-way repeated measures (RM) ANOVAs (between-subject factor: Group; within-subject factor: Time). Tukey’s multiple comparisons was performed as a post-hoc multiple comparisons test, if applicable. For cell counts, one-sample t-tests were used to statistically compare each group’s mean overlap to chance/theoretical mean (1.0). For all tests, alpha was set to p<0.05.

For simple linear regression, X was denoted as the maximum cross-correlation value and Y as the average percent freezing for a single session. The solid line indicated the line of best fit that predicts Y from X with the associated 95% CI. *R*^2^ goodness-of-fit measure, p-value and equation are reported in each plot, as well as in the supplemental Statistical Table.

